# Optimal Placement of High-Channel Visual Prostheses in Human Retinotopic Visual Cortex

**DOI:** 10.1101/2024.03.05.583489

**Authors:** Rick van Hoof, Antonio Lozano, Feng Wang, P. Christiaan Klink, Pieter R. Roelfsema, Rainer Goebel

## Abstract

**Objective:** Recent strides in neurotechnology show potential to restore vision in individuals afflicted with blindness due to early visual pathway damage. As neuroprostheses mature and become available to a larger population, manual placement and evaluation of electrode designs becomes costly and impractical. An automatic method to optimize the implantation process of electrode arrays at large-scale is currently lacking.

**Approach:** Here, we present a comprehensive method to automatically optimize electrode placement for visual prostheses, with the objective of matching pre-defined phosphene distributions. Our approach makes use of retinotopic predictions combined with individual anatomy data to minimize discrepancies between simulated and target phosphene patterns. While demonstrated with a 1000-channel 3D electrode array in V1, our pipeline is versatile, potentially accommodating any electrode design and allowing for design evaluation.

**Main results:** Notably, our results show that individually optimized placements in 362 brain hemispheres outperform average brain solutions, underscoring the significance of anatomical specificity. We further show how virtual implantation of multiple individual brains highlights the challenges of achieving full visual field coverage owing to single electrode constraints, which may be overcome by introducing multiple arrays of electrodes. Including additional surgical considerations, such as intracranial vasculature, in future iterations could refine the optimization process.

**Significance:** Our open-source software streamlines the refinement of surgical procedures and facilitates simulation studies, offering a realistic exploration of electrode configuration possibilities.

## 1 INTRODUCTION

Advances in neurotechnology are revolutionizing the restoration of lost functions through direct brain recording and stimulation (Schalk et al., 2024, Hettick et al., 2022, Sahasrabuddhe et al., 2020, Musk and Neuralink, 2019, Jung et al., 2024, Maynard, Nordhausen, & Normann, 1997). Among these developments, the restoration of a rudimentary form of vision in patients who have become completely blind due to damage in their early visual pathways is particularly relevant (Brindley & Lewin, 1968; Dobelle et al., 1974; Farnum & Pelled, 2020; Nowik et al., 2020; Roelfsema, 2020; Schmidt et al., 1996). The typical approach is to replace a part of the visual pathway with a brain-computer-interface (or visual prosthetic implant) that translates a camera feed into patterned electrical signals that are used to directly stimulate the brain (Normann et al., 2009). Such stimulation of the early visual pathways (retina, optic nerve, thalamus, or visual cortex) elicits dot-like visual sensations with fixed spatial locations known as phosphenes (Brindley & Lewin, 1968; Dobelle et al., 1976; Lee et al. 2000; Maynard 2001; Schmidt et al., 1996). In intact visual systems, the spatial location of a phosphene corresponds to the stimulated neuron’s receptive field, the location of the visual field where visual stimuli evoke a response. Receptive fields are predictably organized to represent the visual world in a way that replicates the topography of the retina across several brain areas (i.e., they have a retinotopic organization). This organization is comparable across individuals, but some variability does exist (Benson et al., 2018a, 2022). Patterns of phosphenes that together form shapes can be evoked by stimulating several electrodes simultaneously (Chen et al., 2020; Dobelle et al., 1976) or in close temporal proximity (Beauchamp et al., 2020; Bosking et al., 2022; Oswalt et al., 2021). This way, phosphene patterns that are created in real-time, based on a (preprocessed) camera feed, will be able to help blind individuals recover some visual functions that were lost or severely impaired after becoming blind (Fernández et al., 2021, Lozano et al., 2020).

The functional properties of artificial vision with different sets of phosphenes, corresponding to different prosthetic devices, can be studied in seeing volunteers using phosphene simulations. To date, such studies have often assumed high density, uniformly spaced, full field covering phosphene configurations (Avraham et al., 2021; Bollen et al., 2019; S. Chen et al., 2009; Sanchez-Garcia et al., 2020; Steveninck et al., 2020; J. Wang et al., 2021), which is a highly unlikely assumption given the anatomy and functional organization of the human brain. Current state-of-the-art prostheses only cover a small portion of the visual field (Fernández et al., 2021; Niketeghad & Pouratian, 2019), which is often a hardware limitation. With these limitations in mind, it is conceivable that different types of daily activities might require different phosphene configurations. A multi-functional visual prosthesis thus requires a careful selection of electrode designs and cortical location target. For example, a dense foveal coverage would be useful for reading, or recognizing an object in front of you, while peripheral vision may be important for context awareness during navigation (Ghezzi, 2023). Additionally, the number and distribution of stimulation electrodes covering visual space are important determinants. More complex phosphene patterns will require a larger number of electrodes and larger patterns require a broader coverage of visual space. Recent developments in image processing have produced deep learning-based algorithms that allow the most relevant features of a scene to be selectively converted into efficient phosphene patterns (Lozano et al., 2020, Beyeler & Sanchez-Garcia, 2022, Van Der Grinten, de Ruyter van Steveninck, Lozano et al., 2024). In theory, visual prostheses should allow crucial every-day life activities such as accurate emotion expression recognition (Bollen et al., 2019), navigation (de Ruyter van Steveninck et al., 2022; Lu et al., 2014; Perez-Yus et al., 2017; Vergnieux et al., 2017; L. Wang et al., 2008), object recognition (Li et al., 2018; Lu et al., 2014; Macé et al., 2015; Sanchez-Garcia et al., 2018; Xia et al., 2015) and even motion detection (Chen et al., 2020; Perez-Yus et al., 2017) to be reestablished after vision loss.

Recent advances in biologically plausible simulations of cortical stimulation-evoked phosphenes allow us to realistically study the functional properties of visual cortex-based phosphene vision (Van Der Grinten, de Ruyter van Steveninck, Lozano et al., 2024), and to simulate and optimize targeted implant visual coverage, as we demonstrate in this work. An ideal phosphene coverage throughout the visual field would likely require many stimulation sites in a brain structure that encodes the entire visual field. However, it remains unclear how phosphenes evoked by stimulation of different hierarchical areas of the visual cortex perceptually combine (Schiller et al., 2011; Schmidt et al., 1996). A major question for all neuroprosthetic approaches is where to interface with the brain. The broad retinotopic organization of the visual brain puts forward many potential targets for implantation, each with their own challenges (Fernández and Normann, 2016). Contemporary approaches therefore typically target a single functional area. Candidate structures are the retina, lateral geniculate nucleus (LGN) and cortical areas V1 (Fernández and Normann, 2016) to higher order regions like V2, V3, V4 and hMT.

Previously, Beyeler et al. (2019) demonstrated computational models of phosphene vision to optimize surgical placement of retinal implants, estimating that up to ∼55% of axonal activation could be avoided, improving the quality of the modeled visual percepts. Bruce and Beyeler (2022) utilized dictionary learning alongside phosphene vision models to enhance expected visual coverage and effective resolution for Argus II users, highlighting the crucial importance of optimization planning in implant design (Beyeler et al., 2017; Granley and Beyeler, 2021). Another candidate structure, the LGN, represents the entire visual space in a relatively small volume of brain tissue, and phosphene perception in non-human primates has been demonstrated by Pezaris and Reid (2007). However, given its location deep in the brain, an LGN visual prosthesis would require long electrode shanks with a high number of densely placed electrodes at the tip.

The primary visual cortex (V1) lies more superficial, making it an interesting candidate for a visual prosthesis. The functional organization of basic visual features such as columnar orientation selectivity and color selectivity is well established in V1 and its position early in the visual processing hierarchy might allow higher level areas in to process more complex visual features, such as motion (Salzman et al., 1990) or faces (Mundel et al., 2003) in a relatively natural way. The feasibility of using large-scale V1 stimulation to generate phosphene-based pattern vision was recently investigated in rhesus monkeys (Chen et al., 2020, 2023) with a 1,024-channel count cortical prosthesis. Human cortex, however, has a lot more gyrification and substantial individual differences in the surface area of visual cortices (Benson et al., 2021). The functional organization of early visual cortex is, however, relatively predictable across individuals as long as the individual anatomy is known (Benson & Winawer, 2018; Rosenke et al., 2021; L. Wang et al., 2015). The ability to derive function from anatomy is especially important in blind patients, as conventional fMRI localizers based on visual input cannot be used. The functional organization of an individual’s brain, together with the electrode design and placement, will ultimately determine what kind of phosphene maps a visual prosthesis can generate.

At this moment, visual prosthetic developments are primarily focused on patients with late-onset blindness because 1) these individuals have experienced vision at some point in their lives and are most aware of their sensory loss, and 2) their visual brain is believed to have developed typically, maintaining the necessary connectivity for these approaches to be effective (Heitmann et al., 2023). In addition to solving scientific, engineering and clinical challenges, it is crucial to not only create technically effective devices but to also align neural implant developers’ perspectives with the needs and expectations of implantees (Nadolskis et al., 2024, van Stuijvenberg et al., 2024). There is, however, a lack of methodologies to predict, optimize and evaluate the impact of electrode design and placement in visual field coverage in large patient populations efficiently.

Our comprehensive and scalable method to optimize the placement of electrodes for a visual prosthesis incorporates 1) the individual anatomy that predicts functional visual maps, 2) flexibility in electrode design, and 3) a preset phosphene map one aims to obtain with the prosthesis. Our pipeline is open-source and automatically finds the electrode configuration that optimally matches a preset ‘ideal’ phosphene map within these constraints. Because the pipeline uses the individual brain anatomy as a starting point, it can also be used in blind subjects after obtaining an anatomical MRI scan. The optimal location and insertion angles of configurable electrode array models are determined with a Bayesian optimization procedure. This algorithm efficiently minimizes a cost function that considers the electrode yield in grey matter, the visual field coverage of the predicted phosphene map, and the similarity in density distribution between the preset target phosphene map and the predicted phosphene map based on the current implantation parameters. We explore the feasibility of an ideal full-field phosphene coverage, and further demonstrate our pipeline by targeting specific subareas of the visual field within a set of practical surgical constraints. In our examples, model parameters and implant location were jointly optimized for a 3D electrode array containing a thousand contact points equally distributed over a 10×10 grid of contact points aimed to be implanted in V1. The general procedure, however, allows for any electrode design and can easily be applied to other or multiple brain areas. We validate our method on 362 human hemispheres using anatomical and retinotopy data from the Human Connectome Project 7T retinotopy dataset (Benson et al., 2018b).

## 2 METHODS

### 2.1 Preprocessing of the Human Connectome Project 7T retinotopy dataset

Data was obtained from the Human Connectome Project 7T retinotopy dataset (Benson et al., 2018). T1-weighted (T1w) and T2-weighted (T2w) structural scans at 0.7-mm isotropic resolution were processed using the FreeSurfer image analysis suite (version 7.2; http://surfer.nmr.harvard.edu). Subject brains were inflated and aligned to FreeSurfer’s anatomical *fsaverage* atlas.

The inferences made in this work are based on the retinotopic data described by Benson et al., (2018). In brief, MRI data was acquired using a Siemens 7T Magnetom actively shielded scanner and a 32-channel receiver coil array with a single channel transmit coil (Nova Medical, Wilmington, MA) at a 1.6mm isotropic resolution and 1s TR (multiband acceleration 5, in-plane acceleration 2, 85 slices). The population receptive field (pRF) maps describe the location and the size of the receptive field for each 1mm isotropic voxel (see Benson et al. (2018) for descriptions of pRF models used). pRF surface maps based on empirical data in Freesurfer fsaverage space were warped to individual space using Freesurfer’s *mri_surf2surf* function. Bayesian inference of the retinotopic maps was performed using Neuropythy’s *register_retinotopy* command (https://github.com/noahbenson/neuropythy). This procedure harmonizes the anatomical inference of the pRFs, the Benson 2014 atlas (Benson et al., 2014) and experimental data (Benson et al., 2018) comprised of retinotopic responses to visual stimuli up to 8 degrees eccentricity. Note that the Benson 2014 atlas predicts maps up to 90 degrees eccentricity, however only the inner 20 degrees of eccentricity in V1, V2, and V3 have been empirically validated.

### 2.2 Pipeline for optimization of electrode grid placement

Our strategy to optimize electrode placement is to minimize the difference between a ‘target’ phosphene distribution (TP) and a ‘predicted’ phosphene map (PP) that we infer from virtual placement of electrodes into a brain model with probabilistic or measured retinotopic maps. This difference was quantified and minimized by calculating the loss with a Bayesian search algorithm (scikit; Pedregosa et al., 2011). An example of the electrode design (and its simplification) used for the simulations is displayed in figure 1A. The main input variables of the optimization function were the angles *alpha* (pitch) and *beta* (yaw), which define the insertion trajectory of the virtually placed implant. This insertion trajectory was determined by finding the intersection between the insertion point on the cortical sheet and the centroid of the calcarine sulcus (CS) at the angles *alpha* and *beta* (Figure 1A-B). The centroid, or geometric center, was calculated using the medians along the three dimensions of the CS volume. Importantly, the CS has been reported to be a reliable estimate of the location and total volume of the human primary visual (Gilissen & Zilles, 1996). To test whether this is also the case in the HCP dataset, the CS volume was determined by Freesurfer’s cortical parcellation algorithm (Desikan-Killiany Atlas) and compared to the volume of V1 (all voxels in the cortical ribbon of the V1-parcel determined by the pRF model), see supplementary figure S1. The range of insertion angles were restricted so that the insertion trajectory cannot intersect with the other hemisphere, and it excludes unrealistic surgical approaches (e.g. insertion in an anterior - posterior direction).

**Figure 1.**
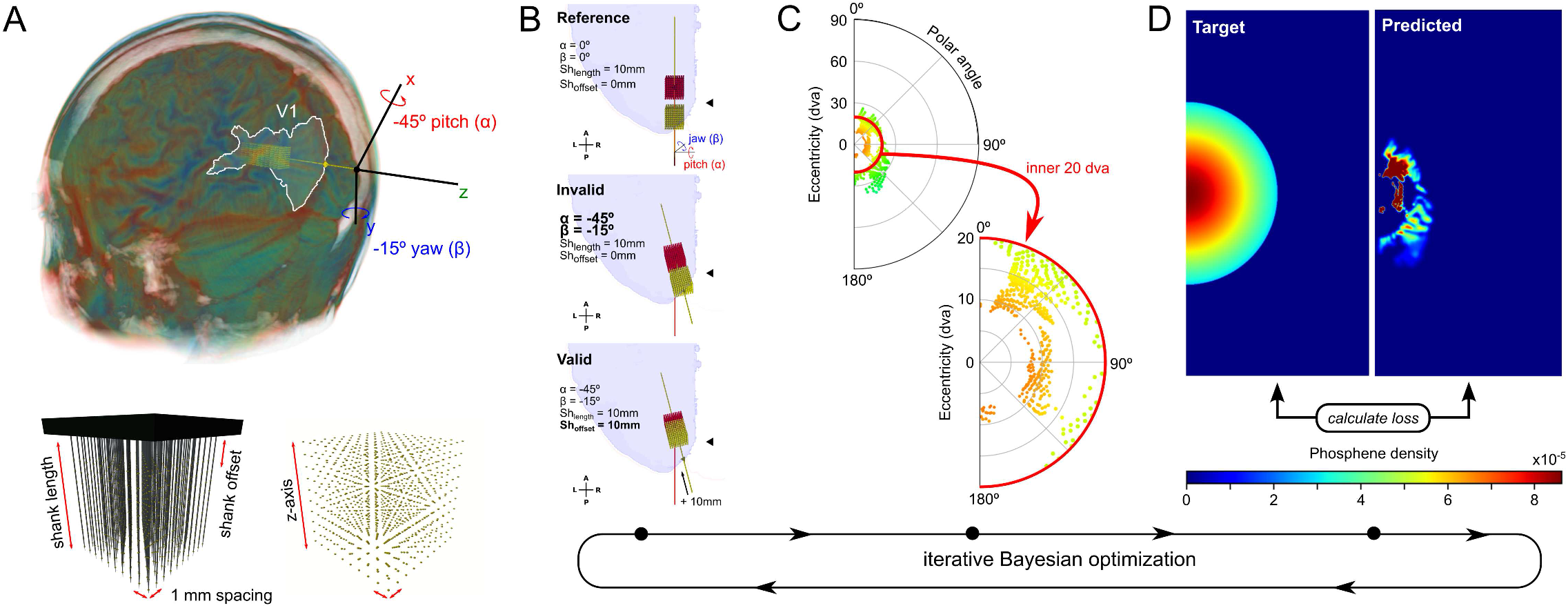
Overview of electrode optimization pipeline. The Bayesian search algorithm determines the next set of parameters based on a loss function with several components (see Methods), and the process is repeated until the optimal set of parameters is found for a specific target phosphene configuration. A) illustration of electrode placement (top) and electrode-grid configuration parameters (bottom). B) The red grid only serves as a reference and is centered on the center of the left calcarine sulcus (black triangles). The implant position (yellow grid) is calculated based on alpha (pitch) and beta (yaw) relative to the reference grid. For a new set of parameters, the resulting configuration can be either valid (left) or invalid (center) when contact points are located outside of the brain. In the ‘valid’ example, the contact point closest to the base is located 10mm from the point where the shank penetrates the cortex. The base is located on the surface of the brain and serves as the anchor point for the shanks. C) Each contact point potentially evokes a phosphene in the polar angle plot. The individual phosphenes are modeled as 2-D Gaussians on a 500-by-1000-pixel phosphene map. Color codes for eccentricity. D) One of the loss terms is the Hellinger distance between the probability distributions of the simulated phosphene map (left) and the target phosphene map (right).

For many combinations of angles, a portion of the electrode grid would end up outside of the cortex. In case any electrode was out of bounds, the configuration was flagged as invalid, and a penalty was assigned in the optimization procedure. An extra parameter (*shank offset)* was added to the optimization function to enable the distance between the insertion point on the cortical surface and the first contact point on the electrode shank to vary (Figure 1A). To further vary cortical depth, the final parameter *shank length* was included in the search for optimal electrode coverage, density and yield (Figure 1A). We constrained electrodes to not exceed a cortical depth of 8cm, as the average width and length of HCP brains are about 17cm and 13cm (Yang et al., 2020). Furthermore, the search can be used to optimally place electrodes in a specific region of interest (ROI). Neurons in V1 have a smaller receptive field size compared to higher visual areas (Benson et al., 2021), potentially allowing denser phosphene maps. Here, we targeted V1 by adding a penalty to the cost function when contact points were located outside V1.

For each set of parameters, or cycle of the optimization pipeline (Figure 2), the electrode-grid is virtually positioned to match the trajectory set by the insertion angles. The predicted phosphene map is then extracted from the intersection of the electrode coordinates and the voxels of the retinotopic map. Each individual electrode that is located in a retinotopically defined map yields a phosphene on a 500-by-1000-pixel image-grid, ranging up to 90 degrees visual angle. Note that the resolution of the image-grid onto which we project the phosphenes impacts the simulation speed. However, if the resolution is too low, the accuracy of the loss function is compromised. To simplify the simulation, we fix the spread of stimulation current to the voxel size of the retinotopic map (1mm^3^). Phosphenes are modeled as 2-D Gaussian spots of light with standard deviation ranging from 0.2 up to 3 visual degrees, depending on the cortical magnification factor and a simulated stimulation amplitude of 100 micro-amps (Tehovnik et al., 2005; Wang et al., 2008). Finally, the process is repeated for a new set of parameters up to 100 times (separately for each hemisphere). The entire procedure is illustrated by the images in figure 1 and the flowchart diagram in figure 2.

**Figure 2.**
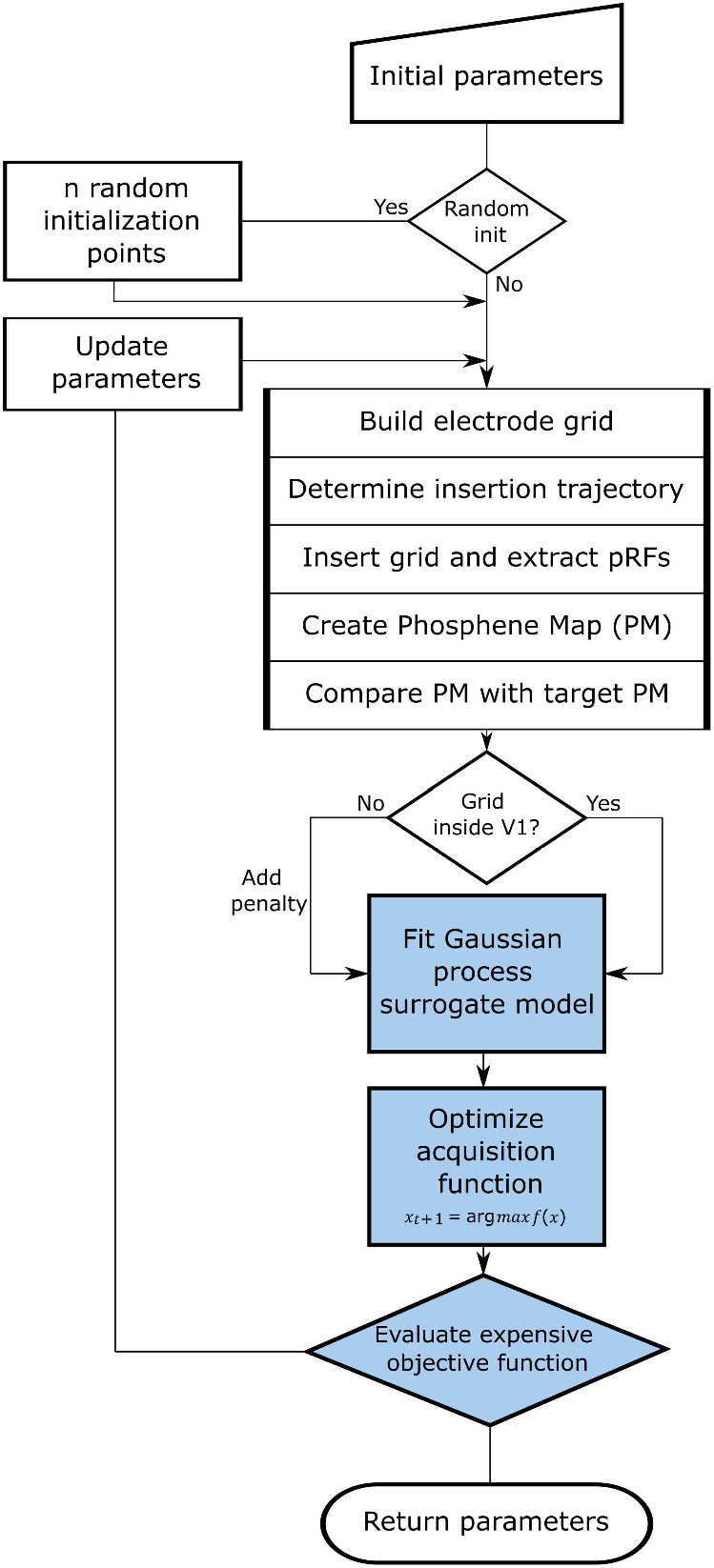
Flowchart diagram Bayesian optimization pipeline. This diagram provides an overview of the virtual electrode placement pipeline. Rectangular boxes indicate processing steps diamond shapes indicate model decisions.

### 2.3 Bayesian optimization

Bayesian optimization has become an attractive method to optimize expensive to evaluate black box functions which are derivative-free and possibly noisy (Shahriari et al., 2016). Bayesian methods have been proven useful in several neurotechnology applications such as neurostimulation interventions (Choinière et al., 2024, Losanno et al., 2021, Bonizzato et al., 2023) and human-in-the-loop optimization of visual encoding in retinal prosthesis (Granley et al., 2023). The algorithm iteratively evaluates a probabilistic model for which a cheap probability function *f* based on the posterior distribution is optimized before sampling the next point. The function objective is considered as a random function (a stochastic process) on which a prior is placed. Here, the prior is defined by a Gaussian process capturing our beliefs about the function behavior. Function evaluations are treated as data and used to update the prior to form the posterior distribution over the objective function. The convergence of the optimization algorithm was accelerated by setting an initial sampling point for which it was known that the resulting grid would hit some portion of V1. We chose to position the center of the electrode grid at the centroid of the calcarine sulcus with initial parameters 0° *alpha*, 0° *beta*, 20mm *shank length* and 25mm *shank offset*. The model can also be run without such prior knowledge of the objective function, but it will likely take more iterations to converge.

### 2.4 Loss function

The loss function comprised three linearly weighted terms that together indicate the difference between a desired phosphene map and the phosphene map predicted from sampling the pRF parameters at the location of the 1000 simulated electrodes in the brain. Below, we explain each loss term and its meaning in the context of the optimization goals. We chose to emphasize the contribution of visual field coverage and phosphene distribution relative to the absolute number of electrodes in V1. However, weights *a, b* and *c* can be tweaked depending on the desired outcome.

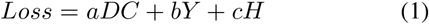

#### 2.4.1 Sørensen–Dice coefficient

The dice coefficient (*DC*) is a measure of overlap between two datasets. DC was computed on the binarized target phosphene map (TP) and a binarized version of the predicted phosphene map (PP). Each pixel was set to 1 if they contain phosphene activation and 0 if they do not. |TP| and |PP| represent the number of elements in each set. The Sørensen index equals twice the number of elements common to both sets divided by the sum of the number of elements in each set. Dice is included to obtain phosphene maps that are localized in the desired visual region.

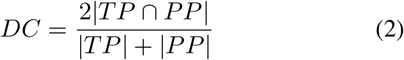

#### 2.4.2 Yield

*Yield* indicates the proportion of electrodes that can evoke a phosphene. We aim this to be as high as possible, thus allowing the implant to achieve high spatial resolution. Contact points outside of the targeted region are penalized by adding a constant to this loss term.

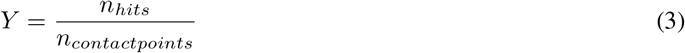

#### 2.4.3 Hellinger distance

The Hellinger distance (*H*) quantifies similarity between two probability distributions and is defined as the square root of the expected squared difference between the square roots of the probability distributions.

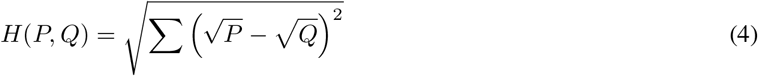

Hellinger distance is especially suitable for comparing normalized discrete probability distributions in machine learning as it is restricted to the range between 0 and 1. *P* refers to the probability distribution of the target phosphenes, and Q to the distribution of predicted phosphenes. The maximum distance 1 is achieved when *P* assigns probability 0 to every set to which *Q* assigns a positive probability, and vice versa. *H* will reward parameter sets for which the virtually implanted electrodes yield a phosphene map with the desired density distributions and penalize density distributions that diverge from the target map.

### 2.5 Phosphene maps

There is a clear (hardware-based) limit to the extent of visual field coverage within the parameters of the single electrode array we use here (evident from Figure 3 and 4). The exclusive consideration of ‘full coverage’ does not provide sufficient insight into the procedure’s accuracy in targeting specific subregions of the visual field. In addition, different cognitive functions may differentially rely on different subregions of the visual field and individual patients may choose a visual field coverage that matches their preferences. Therefore, we demonstrated the spatial specificity of our pipeline by targeting four distinct target coverages where the density of the map increases progressively moving from the periphery towards the center (see Figures 1D, 5 and 6 for visual references):

**Figure 3.**
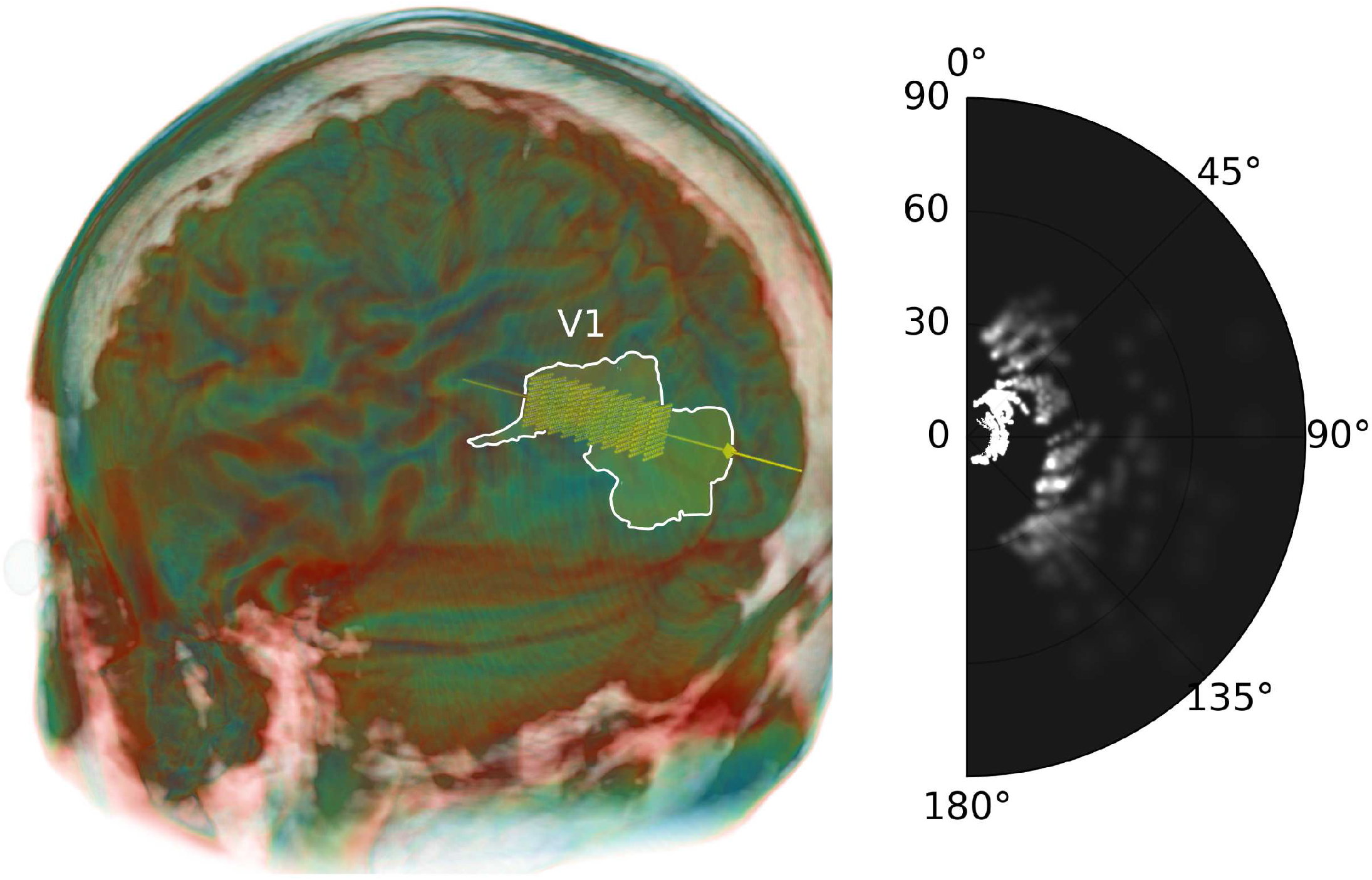
Phosphene configuration in an example subject. On the left the location of the electrodes after implantation are shown for parameters alpha=13, beta=-15, shank offset=13mm, shank length=24mm. VI is indicated as the white ROI. The simulated phosphene map corresponding to the electrode locations is displayed on the right, where intensity depends on the number of overlapping phosphenes.

**Figure 4.**
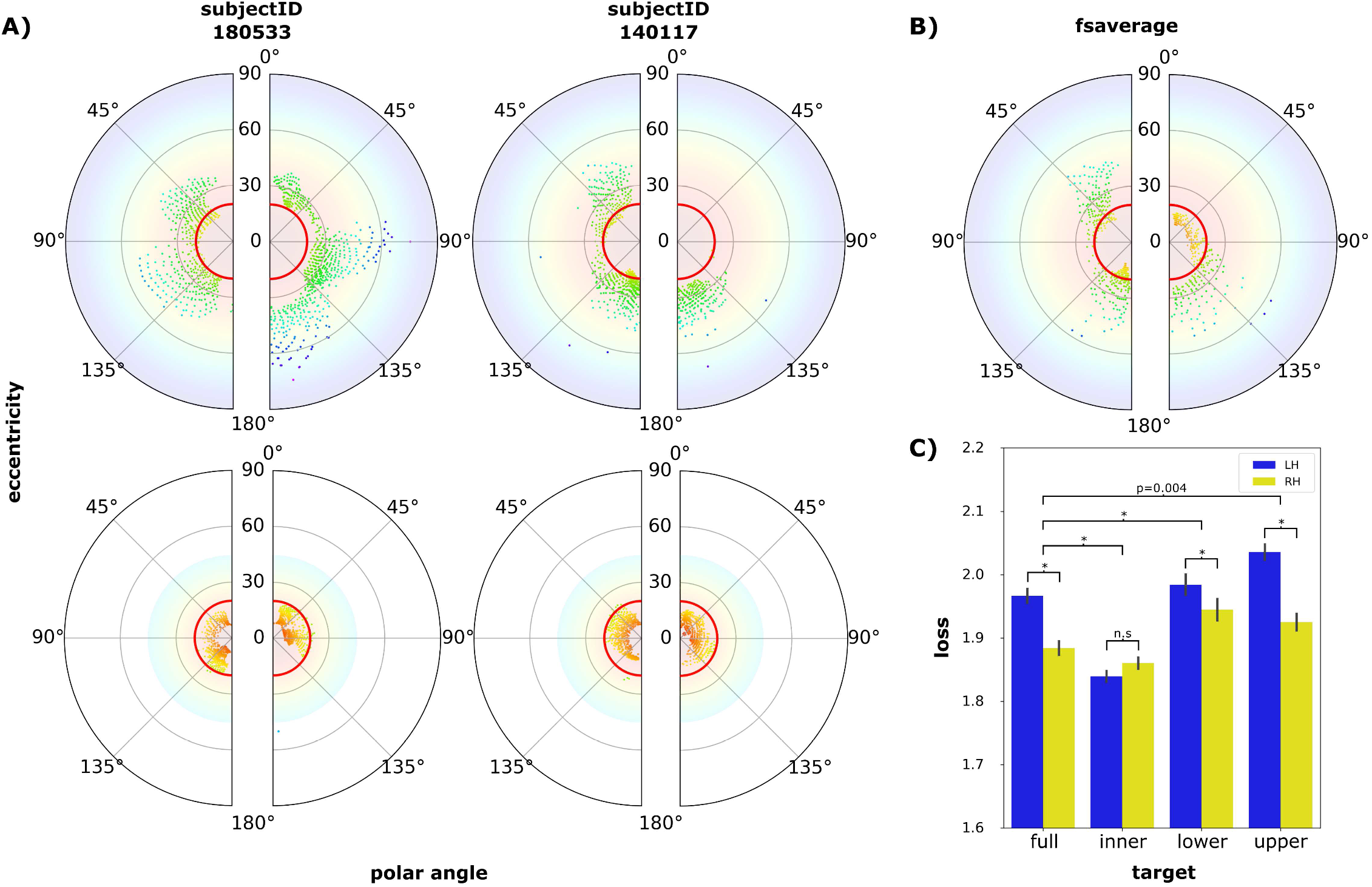
Comparison of individually optimized electrode configurations for the full visual field and foveal target. A) The top row shows phosphene locations in two randomly selected subjects when targeting eccentricities up to 90 degrees (top row) and 45 degrees (bottom row). B) The polar plots show optimization results for the averaged pRF from all subjects in the fsaverage brain (top). The red lines highlight the first 20 degrees eccentricity. C) The barchart (bottom right) depicts loss in all subjects in both hemispheres per target. Error bars indicate standard error. [* = p<0.001].

1. *full target*: covers 0-90° eccentricity.
2. *inner target*: covers 0-45° eccentricity.
3. *upper target*: covers 0-90° eccentricity and 0-45° polar angle.
4. *lower target*: covers 0-90° eccentricity and 135-180 ° polar angle

Simulated phosphenes were scaled based on eccentricity and inspired by recently published phosphene simulation work (Van Der Grinten, de Ruyter van Steveninck, Lozano et al., 2024). Our simulations employed flattened cortical maps as described by Schwartz (1983) and the multi-area visuotopic map complexes in macaque striate and extra-striate cortex identified by Polimeni et al. (2006). Default mapping parameters were taken from Horton and Hoyt’s revision of the classic Holmes map (1991), but patient-specific maps or approximations of Benson et al.’s findings (Benson et al., 2018a) could be used. The cortical magnification factor (CMF) for each electrode location was computed to determine the degrees of visual angle covered per mm of tissue surrounding each electrode. Using a simulated stimulation amplitude of 100 micro-amps, the amount of tissue activated was estimated based on the function and parameters established by Tehovnik and Slocum (2007). CMF was then multiplied by the activated area to obtain the phosphene size in degrees of visual angle. We then drew a Gaussian distribution onto a phosphene center so that 95% of the Gaussian’s activation falls within the phosphene area, with the phosphene diameter equating to 2 standard deviations of the Gaussian.

### 2.6 pRF polar density estimation

Probability density functions (PDFs) of group average PMs were defined by computing a non-parametric kernel-density estimation (KDE) using SciPy 1.0 (Virtanen et al., 2020). The group average PMs were created by summing simulated phosphenes of the optimized electrode locations in all subjects. PMs were created separately for each region of interest (V1, V2, V3, and an all-encompassing ‘grey matter’ ROI). The polar plots in figure 6 allow a qualitative analysis of phosphene locations across subjects. Density was scaled by the maximum to allow for a fair comparison between regions of interest.

### 2.7 Group average electrode configuration

In addition to the individual approach, i.e. determine the best electrode configuration for each person individually, we also optimized electrode placement based on the average brain. Here, we applied the electrode optimization pipeline to the *fsaverage* brain and the group average retinotopy (averaged fMRI timecourse). Phosphene maps were then predicted for individuals either based on the ‘average’ or ‘individual’ parameter estimates. Results were compared to establish the potential benefits of an individual optimization approach (see figure 7).

### 2.8 Statistical analysis

Repeated Measures Individual Analysis of Variance (RMANOVA) and pairwise post-hoc Tukey’s multiple comparison tests were performed using Python library statsmodels v0.14.0 (Seabold & Perktold, 2010). The reported statistical results were Bonferroni corrected by dividing the significance levels by two (the number of individual ANOVA analyses) to correct for multiple comparisons.

## 3 RESULTS

Optimal electrode placement in primary visual cortex was determined for both hemispheres of 181 subjects from the HCP 7T dataset by minimizing the difference between a desired phosphene map (DPM) and predicted phosphenes from virtual electrode placements. The feasibility of obtaining maximal visual field coverage is illustrated using data from example subjects (figure 3 and 4). The average results over all individuals are then described using pRF/phosphene density distributions (figure 5 and 6) indicating the probability that a phosphene was evoked in a specific spatial location with the optimized placements parameters. Finally, we compared individualized optimization based on an individual’s own brain scan with the general optimization solution based on the group (fs)average brain (figure 7).

**Figure 5.**
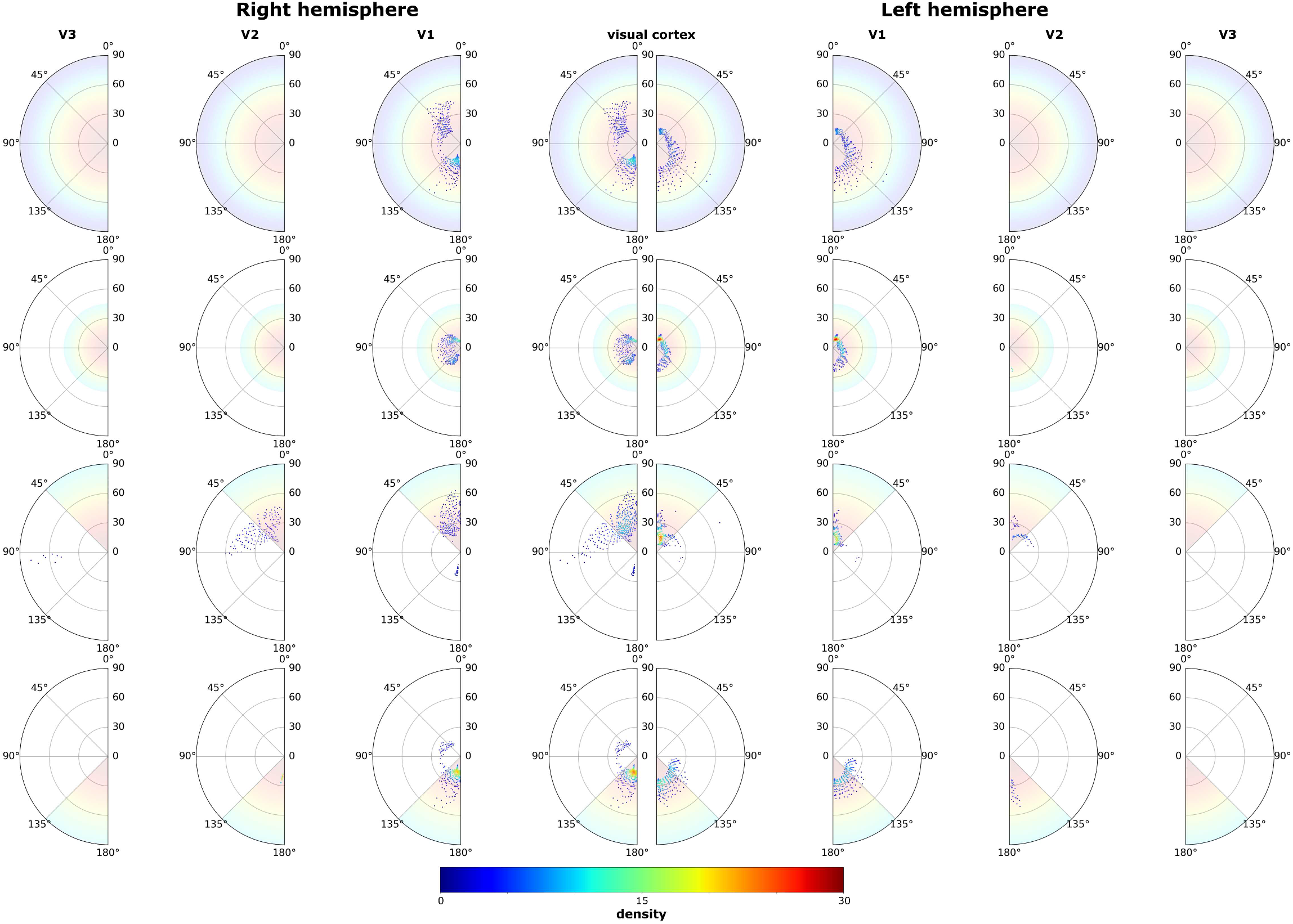
pRF density polar plots in the fsaverage brain. Results after running the pipeline on subject average pRF maps projected on the fsaverage brain. The color indicates the density (as determined by Gaussian KDE) of simulated phosphenes in visual space. For reference, the target PM is shown on the background of each plot. For the lower and upper target, even the best solution places a notable amount of electrodes in V2.

**Figure 6.**
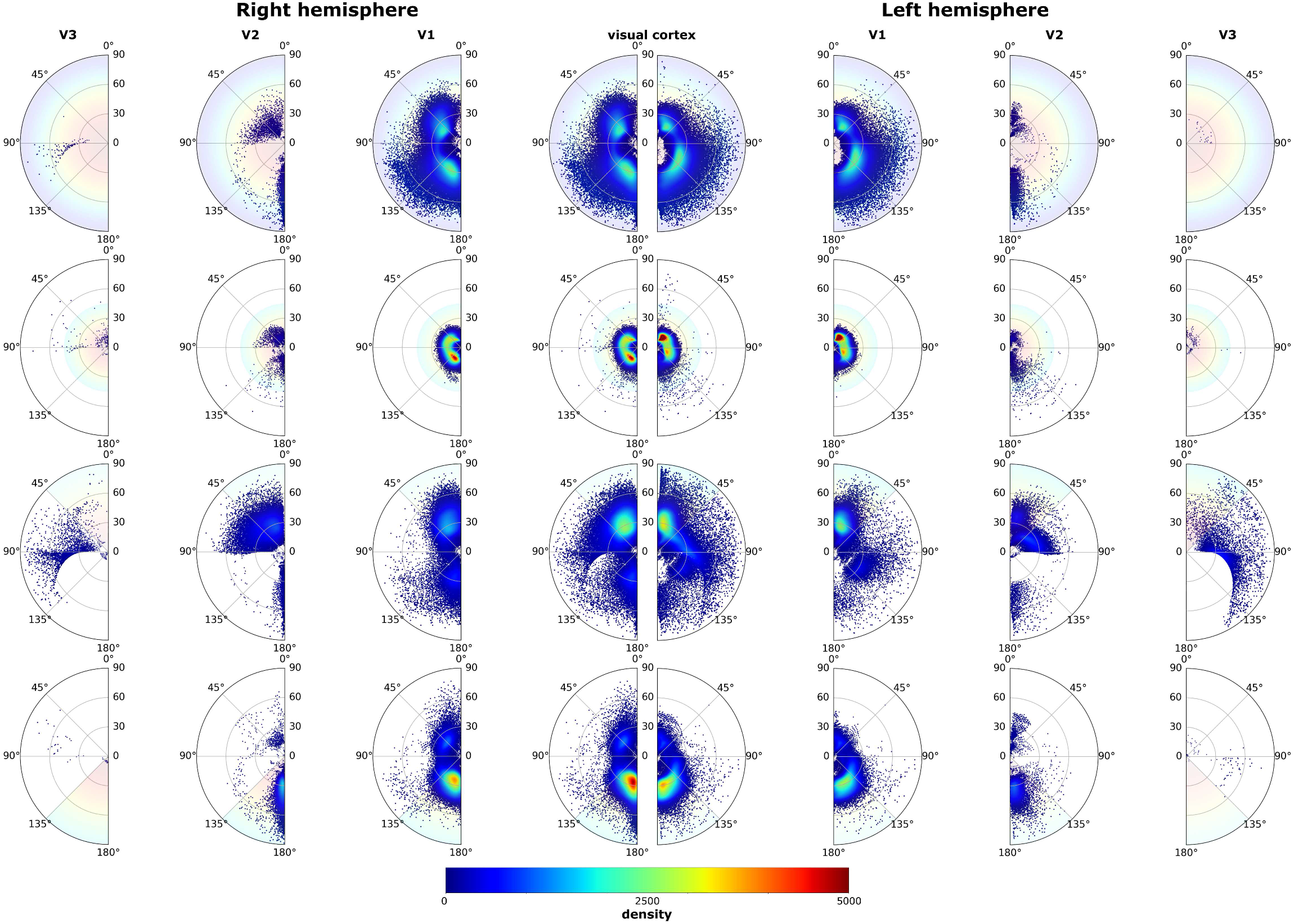
Group average pRF density polar plots. The center plots show a summary of all pRF locations in either left or right hemisphere after optimizing electrode configurations in all subjects. The color indicates the density (as determined by Gaussian KDE) of simulated phosphenes in visual space. As a reference, the target PM is shown on the background of each plot. For many individuals and depending on the target visual field region, many electrodes are placed in V2 and V3.

**Figure 7.**
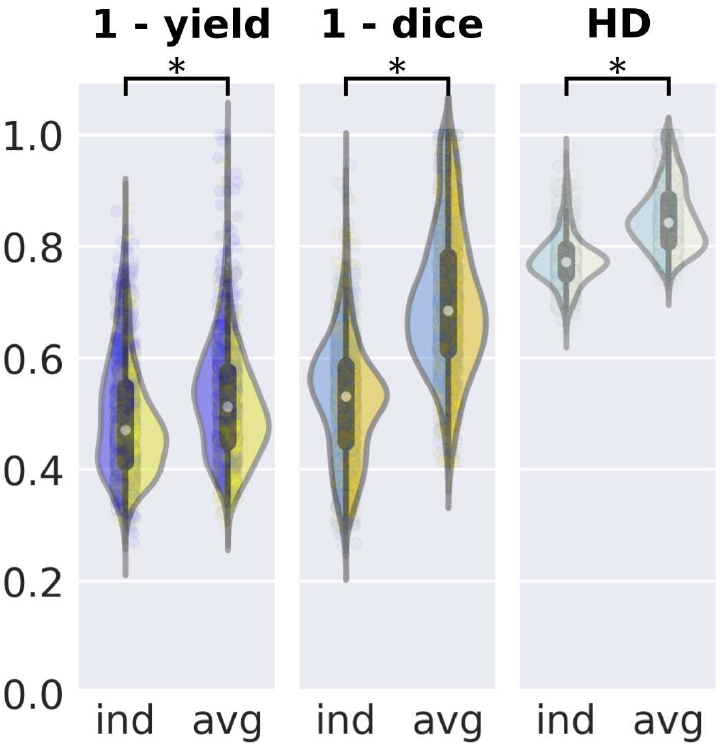
Superior visual coverage for individually fine-tuned electrode placement. The violin plots describe the three loss terms with each half representing results for the left or right hemisphere. Configurations based on individually optimized parameters (ind) result in significantly more electrodes successfully placed in V1, higher phosphene coverage and a more accurate density distribution compared to an average electrode configuration (avg) [* = p<0.001].

### 3.1 Standardized starting point for the Bayesian search space

In our sample of 181 individuals, we found that the volume of the calcarine sulcus (CS) correlated well with the volume of V1 in both hemispheres (Pearson correlation left hemisphere: *r* = 0.95, p < 0.001; right hemisphere *r* = 0.87, p < 0.001; figure S1). We used the centroid of the CS as a standardized anatomy based starting point for optimization (with alpha and beta insertion angles set to zero).

### 3.2 Phosphene maps from optimized electrode placements

We optimized electrode placements for a target phosphene density map with a coverage of 90 degrees and with a density that follows a Gaussian function, corresponding to the entire visual field. Figure 3 provides an example of the best possible placement for our electrode array design, using the trajectory constraints defined in ‘Pipeline for optimization of electrode placement’, the brain’s anatomical dimensions, as well as the predicted phosphene density map. The optimal phosphene map reveals a densely populated area between the first 5 to 20 visual degrees, followed by a coherent spaced-out distribution at higher eccentricities. High eccentricities are notably absent, due to constraints of the electrode array and cortical layout.

Figure 4 shows optimized phosphene locations in two subjects for the full visual field and the inner half of the visual field, along with the best configuration for the *fsaverage* brain. Repeated measures ANOVA revealed a significant interaction on *loss* between *target* and *hemisphere* (F(3, 540) = 7.1, p < 0.001). We also found a pronounced difference between *left* and *right* hemispheres for *full* target (M = -0.05, p < 0.001), *lower* target (M =-0.10, p < 0.001) and *upper* target (M = -0.07, p < 0.001). Loss was lower for the right hemisphere for all targets, except the inner target (see Figure 4C).

Optimization variability across the population was further evaluated with group-average pRF density functions of predicted phosphenes across the visual field. In figure 5 and 6, pRF density plots are shown per target phosphene map across the regions of interest. We observed a fairly symmetrical phosphene peak distribution across hemispheres, with most phosphenes located quite central in the visual field. Phosphenes obtained for the upper and lower target maps were mostly located in the expected visual quadrants, however a relatively large proportion of these phosphenes came from electrode contacts in V2 and V3. This indicates that even when the algorithm tends to place the grid in V1 for these target regions, frequently no solution could be found where all 1000 electrodes are placed in V1.

### 3.3 Group average optimization versus individual optimization

For a less computationally expensive, and thus faster way to design an electrode grid, it is possible to optimize the placement of electrodes on the average brain and then virtually implant this configuration in all individuals. In figure 7 this group average approach is compared with the results from the individualized optimization approach. Each element of the loss function is shown separately for a more detailed description of differences in visual coverage. Post hoc comparisons using the Tukey HSD test indicated that the individually optimized approach significantly outperformed the average approach for phosphene yield (M = 0.03, p < 0.001), phosphene map coverage (M = 0.18, p < 0.001), as well as Hellinger distance (M = 0.08, p < 0.001). See supplementary materials (figure S3) for single-subject examples.

## 4 DISCUSSION

Restoring a rudimentary form of vision in blind patients is an ambitious aim for neuroscience and neural engineering that has recently gained a lot of attention. Visual prostheses that interface directly with the brain may one day be a realistic treatment option for blindness that results from damage to the early visual pathways Normann et al., 2009; Fernández et al., 2021; Roelfsema, 2020; Rosenfeld et al., 2020; Stephen Shankland, 2022). However, the development of such a treatment comes with significant challenges. The method of optimizing electrode placement that we present here addresses several of these, allowing to optimize the design and placement of electrode arrays to obtain a functional implant with coverage and resolution that matches a pre-defined target, within predetermined boundaries for surgical implantation.

Our data-driven solution uses simulated phosphenes based on individual anatomy, and efficiently optimizes electrode placement, minimizing the difference between the simulated phosphene patterns and a predetermined target layout. While the general method works for arbitrary electrode array designs, we here demonstrate it using a 1000-channel 3D electrode array. Even though we set the algorithm to optimize electrode placement in V1, a fraction of the contact points ended up in V2 and V3. The algorithm also has the capacity of optimizing placement based on multiple visual areas, while it is currently unknown how phosphenes that are evoked in distinct visual areas are perceptually combined into patterns. Future studies should address this question, as it would greatly expand the possible configurations for a cortical visual prosthesis.

The predicted phosphene maps were generally closer to the target distribution for the right hemisphere for all targets, except the inner target. This difference might be related to anatomical hemispheric differences (see also supplementary figure S1). Our results also demonstrated that while it is possible to base electrode placement on the optimization solution in the average brain, an individually optimized approach performed significantly better. This finding underscores the importance of tailoring electrode placement and design to the specific anatomical characteristics of each individual (Beyeler et. al, 2019, Bruce and Beyeler, 2022). By considering individual variations, we can achieve better outcomes and optimize visual restoration for each patient.

Covering the entire visual field with high spatial resolution is unfeasible with currently available electrodes such as the type of single rigid electrode grid we simulate here. It might be achieved by distributing electrodes over multiple (smaller) electrode arrays, but challenges emerge when trying to optimize the placement of multiple interdependent arrays. For example, they cannot physically occupy the same space, and their implantation trajectories need to be compatible. Our current approach can be expanded to place multiple electrode arrays in a serial manner, however more elaborate optimization procedures may be needed. About half of the electrodes in our simulations were located inside grey matter, with most of the remaining electrodes likely located in white matter. Here, we assumed that white matter stimulation cannot effectively create useful phosphene percepts. This assumption is partly based on the lack of a clear prediction for the retinotopic organization of white matter. However, since these fiber paths do connect visual areas and carry visual information, further research might help to understand the perceptual effects of white matter stimulation.

The loss terms implemented in our approach were chosen for interpretability and ease of use. However, they can easily be customized, replaced, or expanded, which makes the implant optimization framework rather flexible. With the example electrode array used here, array placement results were more successful for the foveal (inner) target region, compared to the more peripheral (full, upper and lower) target regions. This is a direct consequence of the higher eccentricities being located deeper along the calcarine and requiring longer shanks to reach them, which automatically increases the electrode spacing along the shank. In addition, a relatively large part of cortex is dedicated to central vision (low eccentricities) compared to peripheral vision (known as cortical magnification). A more suitable approach to get coverage from the peripheral visual field is to target different sections of the visual field with multiple electrode arrays and/or in different visual areas. Retinotopic maps are ubiquitous along the visual cortical hierarchy (Wandell et al., 2007), but receptive field sizes increase towards higher areas, reducing the spatial resolution of potential phosphene vision. Importantly, and to reiterate a previous point, it is currently unknown how simultaneous stimulation across hierarchically different visual areas perceptually combines.

The accuracy of the phosphene map predictions in our pipeline crucially depends on 1) the precision and the spatial resolution of the underlying population receptive field maps/atlas. There is an accumulative uncertainty that stems from alignment and extrapolating data from seeing individuals to blind individuals. 2) the assumptions of the phosphene simulation algorithm. A possible improvement to the first matter could be to acquire higher quality pRF data. The availability of higher field strengths could potentially yield scans with higher resolution. It is highly recommended to obtain detailed individual anatomy using a high-resolution MRI scan whenever possible. Retinotopy can then be estimated based on individual anatomy and group-average (probabilistic) pRF maps, yielding the best possible results. Finally, the current version of our pipeline does not account for intracranial vasculature, which is another surgical constraint for safe implantation. Future versions of the pipeline might be expanded to minimize the amount of brain tissue traversed to reach the target location, and include hard constraints to avoid large vasculature. The latter would require more detailed structural scans of individual patients’ brains.

In conclusion, we present an automated approach to optimize the electrode placement in human visual cortex for visual neuroprostheses. It efficiently finds a suitable location yielding the closest possible match to a preset visual field coverage. We demonstrate its potential for large-scale use on a dataset comprising of anatomical and pRF data for 181 individual brains (362 hemispheres). Our open-source software (https://github.com/rickvanhoof/vimplant) can easily be expanded to facilitate different electrode configurations (including multiple arrays), different (multiple) target areas, or constraints (through different cost functions). As such, it will be able to refine surgical procedures as well as drive simulation studies in healthy observers with phosphene vision configurations that are realistic representations of the possibilities with any given electrode design.

## Acknowledgements

This work was supported by the Dutch Organization for Scientific Research (NWO): STW grant number P15-42 ‘NESTOR’, ALW grant number 823-02-010 and Cross-over grant number 17619 ‘INTENSE’. This work was carried out on the Dutch national e-infrastructure with the support of SURF Cooperative.

