## supplemental material for "Optimal Placement of High-Channel Visual Prostheses in Human Retinotopic Visual Cortex"

### SUPPLEMENTARY MATERIALS

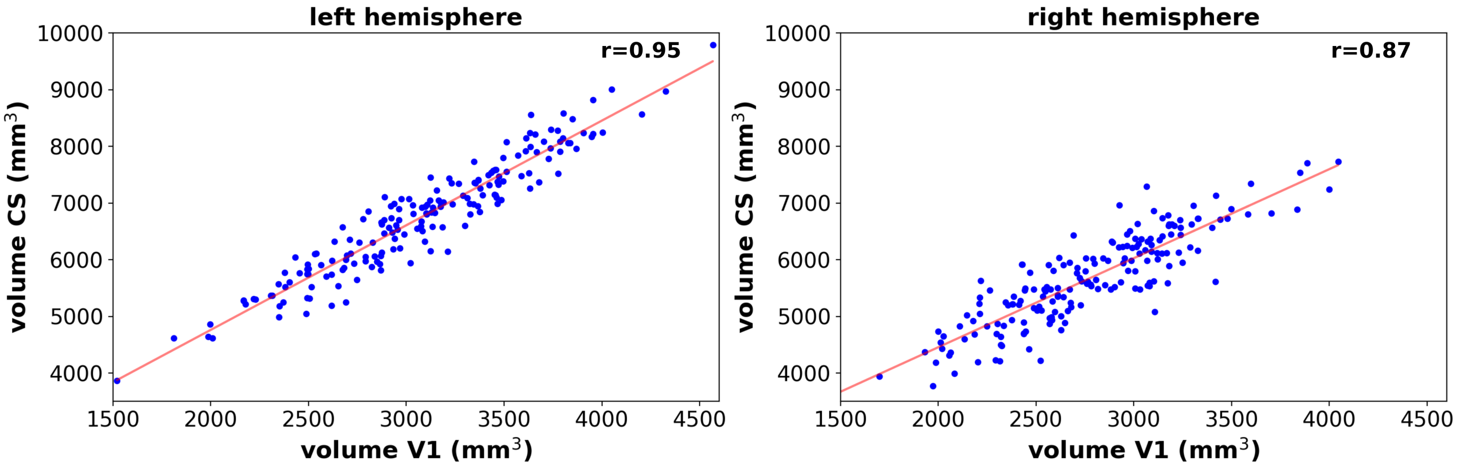


**Figure S1.** Volume comparison between CS and V1 for the left (left) and right (right) hemisphere ($p < 0.001$).

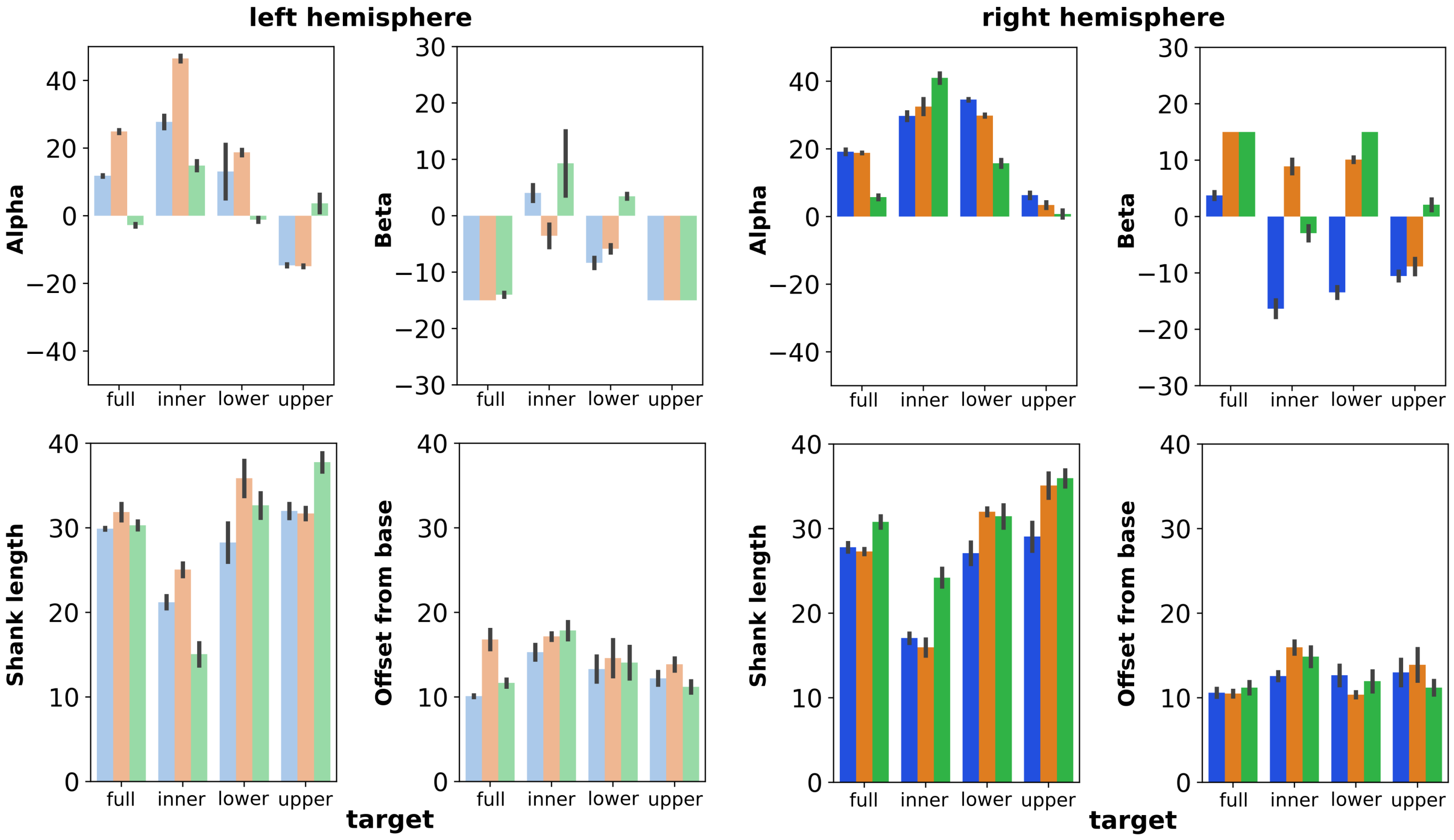


**Figure S2.** Within-subject reliability. The figure shows mean electrode placement parameters within three randomly selected individuals (with standard error bars) after optimization was repeated 20 times. The error bar is an indication of within-subject variability and is relatively stable compared to between-subject variability.

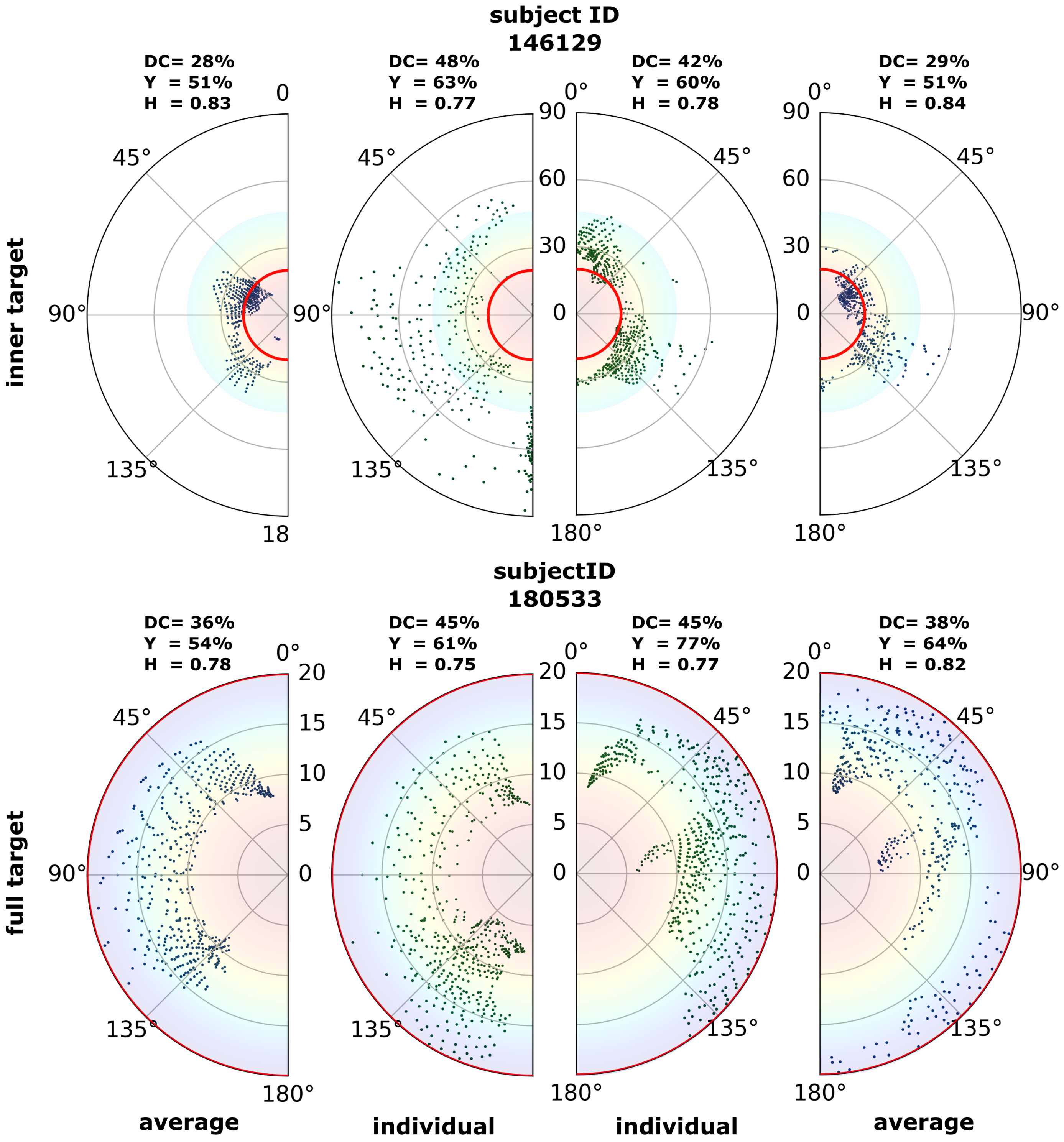


**Figure S3.** Phosphene locations for two example subjects comparing individual versus group-based optimized parameters.
